## Supplemental for "The transcriptional response of cortical neurons to concussion reveals divergent fates after injury"

### Supplemental Figure 1

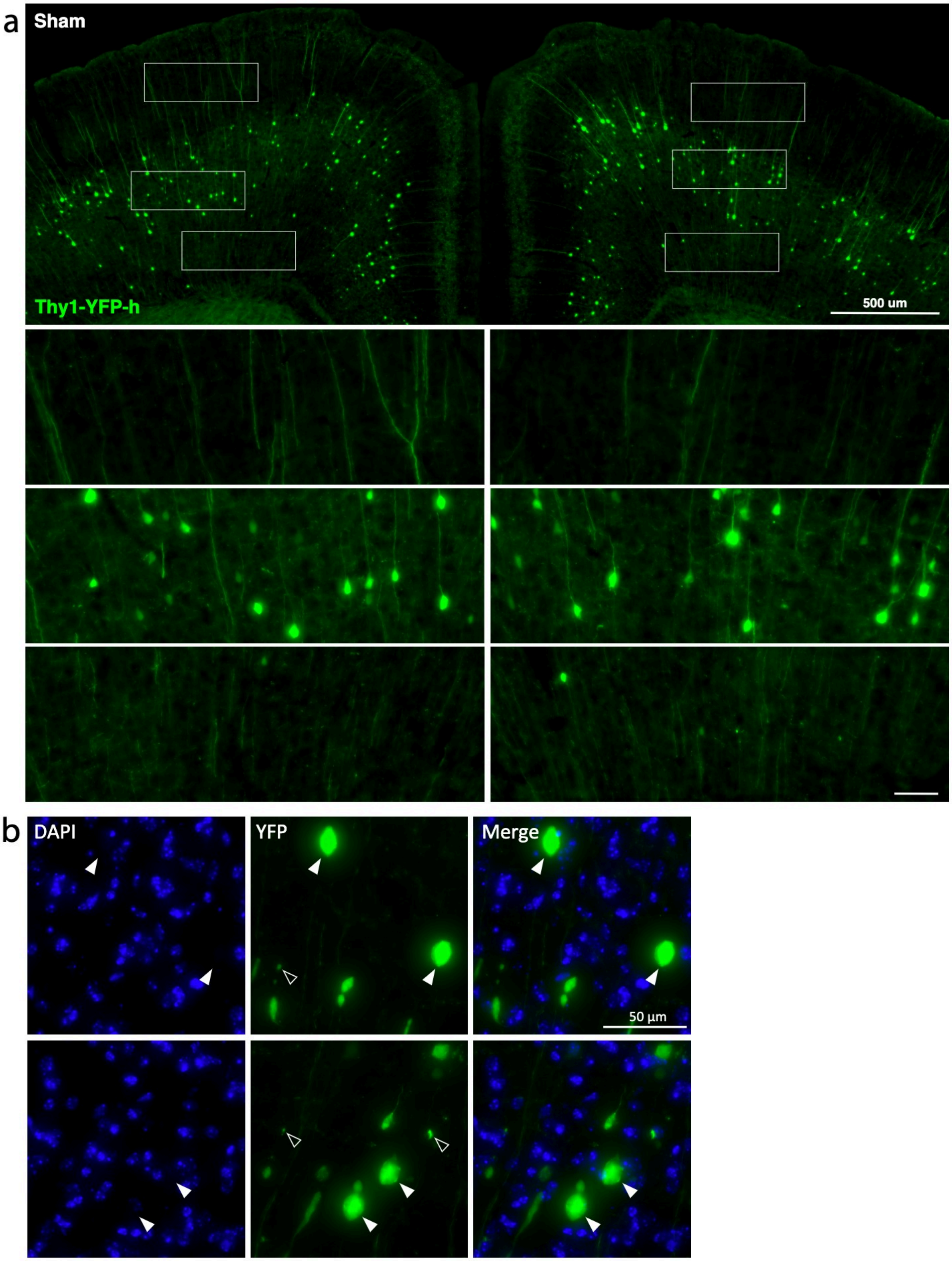

**Supplemental Figure 1. Thy1-YFP expression in sham and classification of axon blebs. a.** Example coronal section from an uninjured sham Thy1-YFP animal with dendrites, cell bodies, and axons highlighted below. Scale bar representative for all insets. **b.** High magnification images showing DAPI expression and the axons of Thy1-YFP animals after mTBI. Closed arrows highlight DAPI-negative axonal swellings, open arrows highlight axon beading. Lack of DAPI-staining was an inclusion criterion for axon swellings. Low magnification scale bar, 500 µm. High magnification scale bar, 50 µm.

### Supplemental Figure 2

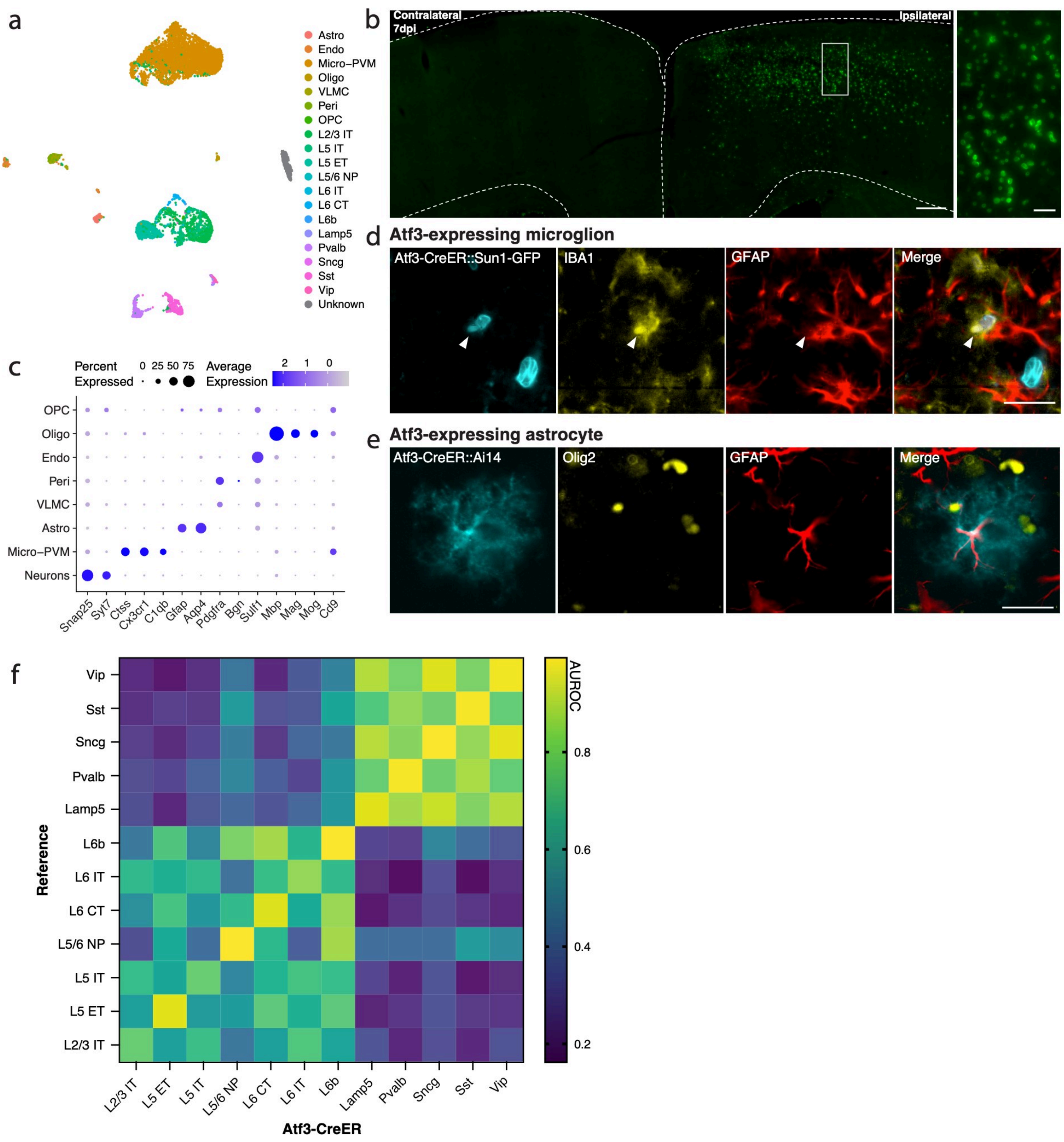

**Supplemental Figure 2. Validation of Atf3-expressing cell types. a.** UMAP of all *Atf3*-expressing cells collected at 7 dpi from the Atf3-CreER mouse, annotated by mapping to a reference atlas. One neuronal cluster could not be confirmed as cortical, labeled as “Unknown”. **b.** Coronal section of cortex showing expression of Atf3-CreER::Sun1-GFP in mice used for sequencing (tamoxifen at 4 and 5 dpi, tissue collected at 7 dpi). Inset highlights large and small nuclei in cortical layers V and II/III. **c.** Dotplot of general cell type markers confirming the reference mapping shown in a. **d.** Validation of the existence of Atf3-expressing microglia seen in the sequencing. Atf3-CreER::Sun1-GFP signal highlights an Atf3-expressing nucleus (cyan) expressing IBA1 (yellow), but not GFAP (red). **e.** Validation of the existence of Atf3-expressing astrocytes seen in the sequencing. Atf3-CreER::Ai14 signal highlights an Atf3-expressing astrocyte (cyan) expressing GFAP (red), but not Olig2 (yellow). **f.** Heatmap of AUROC scores from MetaNeighbor between neurons in the reference dataset (y axis) and neurons in the Atf3-CreER dataset (x axis). Legend indicates dark blue is a low AUROC, meaning low similarity between datasets, and yellow is a high AUROC, meaning high similarity between datasets. Low magnification scale bar, 200 µm. High magnification scale bar, 25 µm.

### Supplemental Figure 3

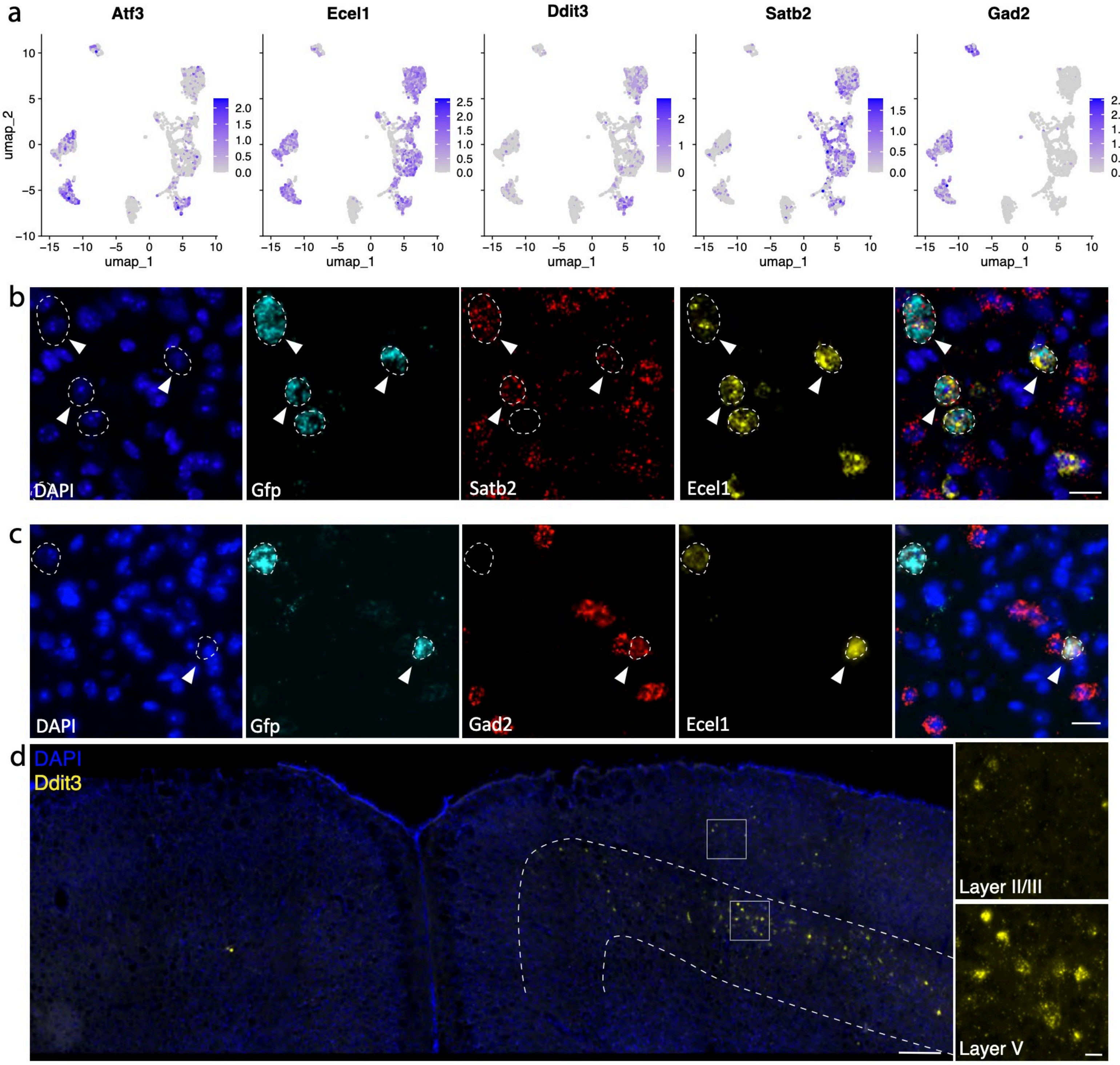

**Supplemental Figure 3. Validation of Atf3-expressing neuron types. a.** Feature plots of *Atf3, Ecel1, Ddit3* and markers for Atf3-expressing neuron groups; *Satb2* for injured excitatory neurons, *Gad2* for inhibitory neurons. **b.** ISH highlighting injured excitatory neurons, showing coexpression of *Ecel1* (yellow) and *Satb2* (red) in *Gfp*+ cells (cyan) in the Atf3-CreER::Sun1-GFP injured cortex at 7 dpi. *Gfp*+ cells are outlined. Arrowheads pointing at *Gfp*+ injured excitatory neurons. **c.** ISH highlighting injured inhibitory neurons, showing coexpression of *Ecel1* (yellow) and *Gad2* (red) in *Gfp*+ cells (cyan) in the Atf3-CreER::Sun1-GFP injured cortex at 7 dpi. *Gfp*+ cells are outlined. Arrowhead points to a *Gfp*+ injured inhibitory neuron. **d.** ISH of *Ddit3* (yellow) shown in ipsilateral and contralateral cortices. Layer V is outlined. Insets highlight ipsilateral neurons in layer II/III (top) and layer V (bottom). Scale bars, 20 µm.

### Supplemental Figure 4

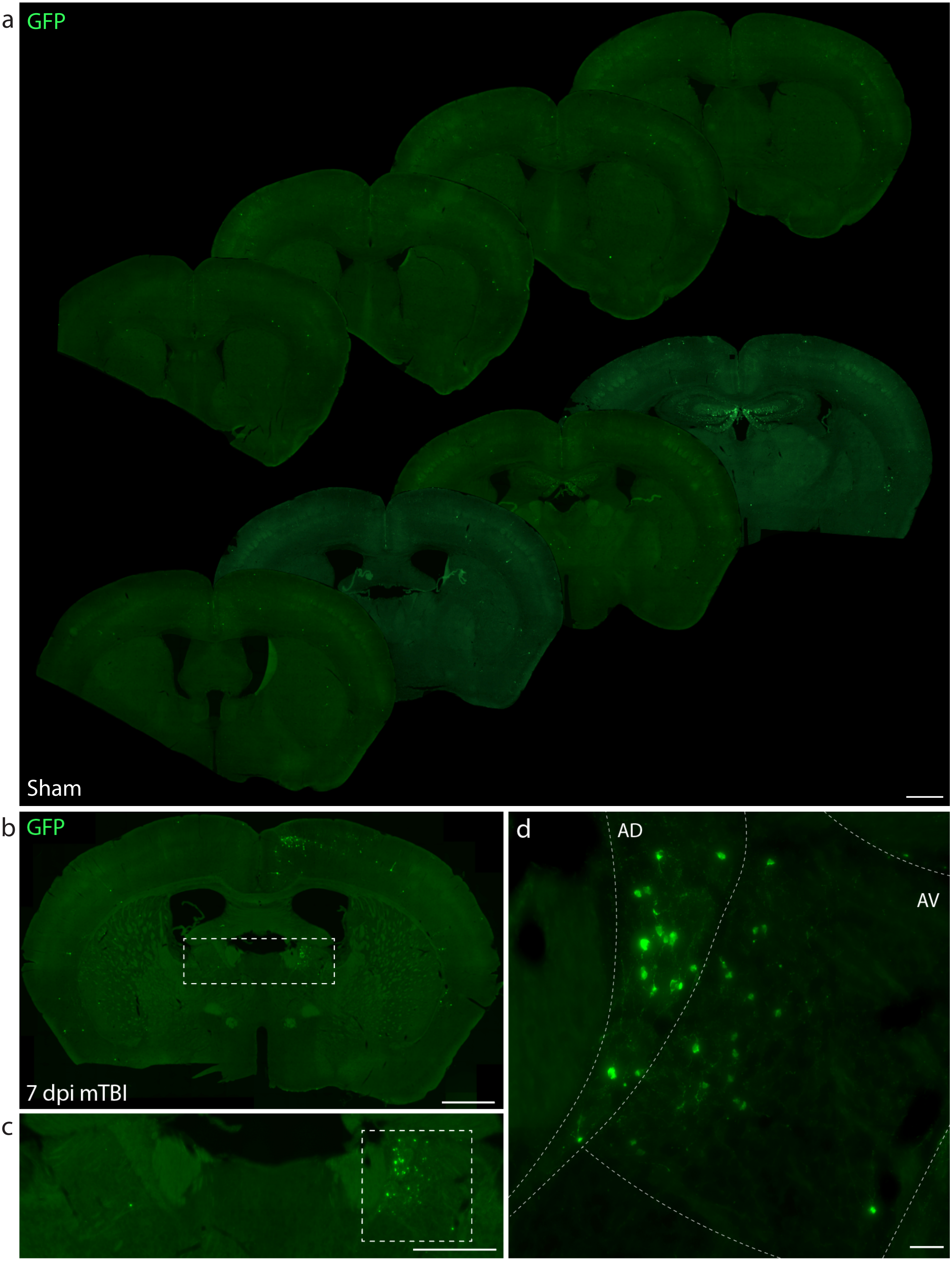

**Supplemental Figure 4. Atf3-Cre::Snap25-GFP expression at 7 dpi. a.** Coronal sections from Atf3-Cre::Snap25-GFP sham animals (uninjured) showing immunolabeling of GFP without injury. Many hippocampal neurons express GFP, likely due to developmental Atf3 expression. The right ventral portion of the brain was notched. **b.** Image showing a section with GFP immunolabeling at 7 dpi in the ipsilateral cortex and ipsilateral anterior thalamic nuclei. **c.** Inset shows bilateral anterior thalamic nuclei, with GFP immunolabeling in the ipsilateral nuclei. **d.** Inset of the ipsilateral anterior thalamic nuclei. The anterodorsal (AD) and anteroventral (AV) thalamic nuclei are outlined. Low magnification scale bars, 1 mm. Scale bars in **b** and **c**, 500 µm. Scale bar in **d**, 50 µm.

### Supplemental Figure 5

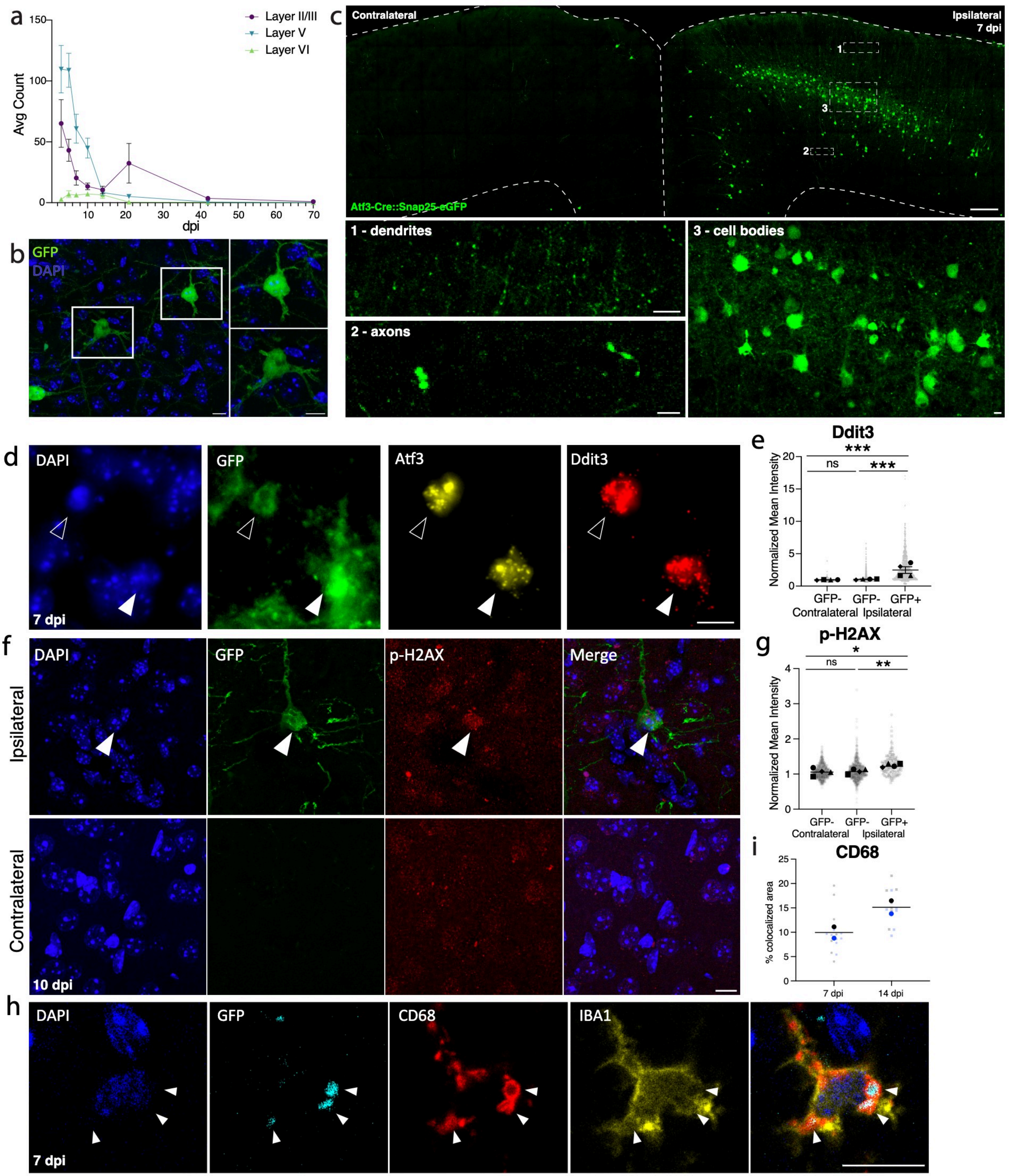

**Supplemental Figure 5. Layer V Atf3-GFP neurons undergo apoptosis by 14 dpi. a.** Quantification of ATF3 protein expression (regardless of cell type) by cortical layer at 3, 5, 7, 10, 14, 21, 42, and 70 dpi. N=6 per timepoint, 3-4 sections counted per animal. Error bars represent SEM. **b.** High magnification image of layer V GFP-expressing neurons highlighting a neuron that appears morphologically healthy (top) and one that appears unhealthy (bottom). **c.** Low magnification image of ipsilateral and contralateral cortex in Atf3-Cre::Snap25-eGFP mice. High magnification images of dendrite region (1) highlighting degeneration, axon region (2) highlighting axon swellings, and cell body region (3) highlighting variability in neuron morphology. **d.** In situ hybridization of Atf3 and Ddit3 in Atf3-GFP tissue highlighting layer V GFP+ neurons that appear morphologically unhealthy and co-express these genes. Open arrowhead highlights neuron with condensed DAPI, closed arrowhead highlights neuron with low GFP expression. **e.** Quantification of mean intensity of Ddit3 mRNA in layer V GFP+ neurons and GFP- neurons of the ipsilateral and contralateral cortices at 7 dpi normalized to background intensity. Average per animal and value per cell are displayed. Each shape represents one animal (*** $p < 0.0001$ by linear Mixed Effect modeling allowing for different variances per group). **f.** Immunostaining of phospho-H2AX in Atf3-GFP tissue in ipsilateral (top) and contralateral (bottom) cortex. Arrow highlights a layer V GFP+ neuron that appears vacuolized and expresses high levels of phospho-H2AX. **g.** Quantification of mean intensity of phospho-H2AX in the nuclei of GFP+ neurons and GFP- neurons in the ipsilateral and contralateral cortices in layer V at 7 dpi normalized to background intensity. Average per animal and value per cell are displayed. Each shape represents one animal (* $p = 0.0219$, ** $p = 0.0069$ by linear Mixed Effect modeling allowing for different variances per group). **h.** High magnification of a microglial cell (IBA1, yellow) at 7 dpi. Arrows highlight microglial lysosomes (CD68+, red) engulfing GFP+ debris (cyan). **i.** Quantification of percent of CD68+ area colocalized with GFP, reflecting CD68 engulfment of GFP+ debris at 7 and 14 dpi. Average per animal and value per ROI are displayed. Each color represents one animal. Scale bars, 10 µm.

### Supplemental Figure 6

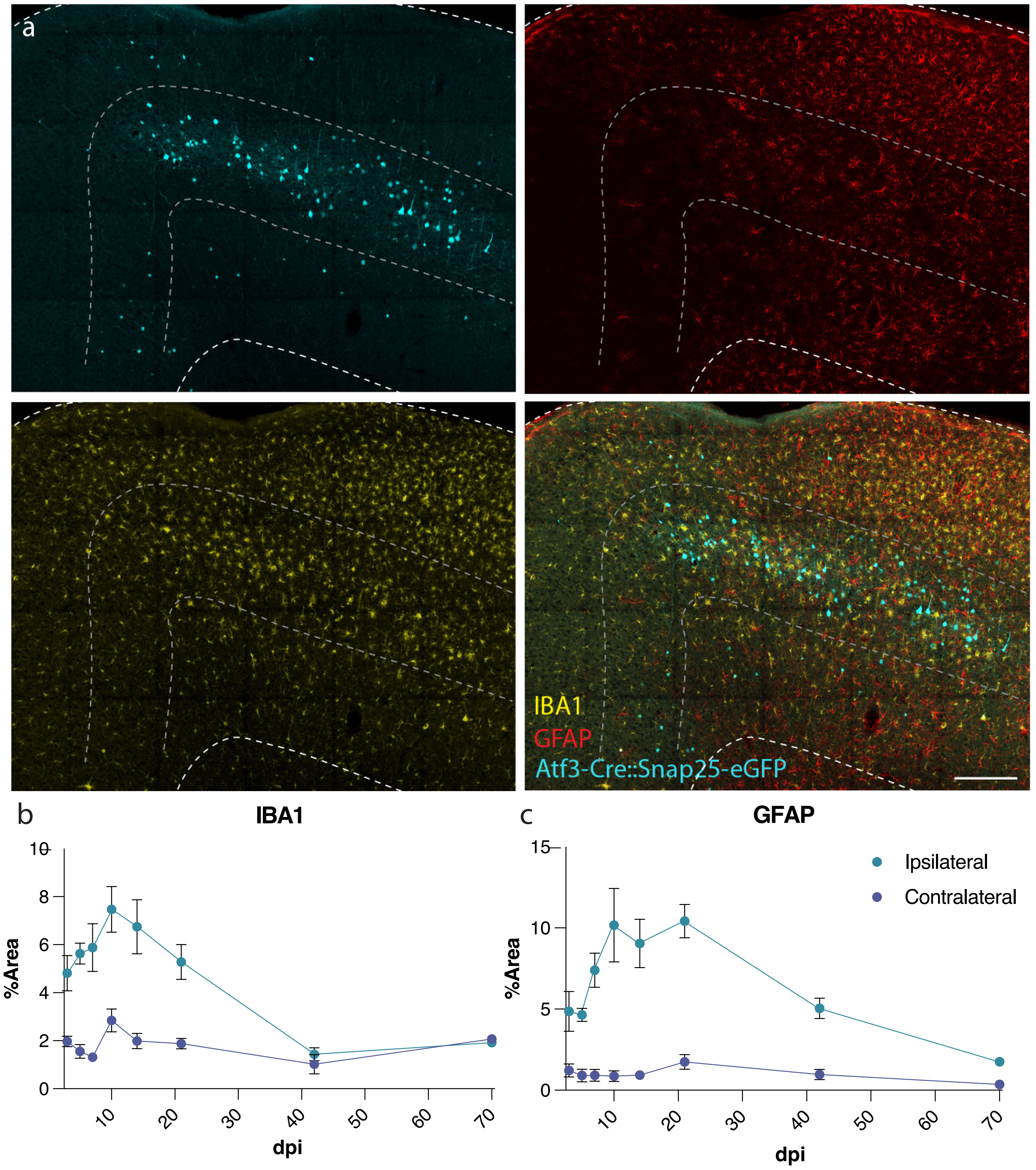

**Supplemental Figure 6. Neuroinflammation timecourse following mTBI. a.** Immunostaining for IBA1 (yellow) and GFAP (red) in ipsilateral cortex of Atf3-Cre::Snap25-eGFP tissue at 7 dpi, highlighting the double layer pattern of microgliosis around GFP+ neurons (cyan) and the injury site and the extent of astrogliosis throughout the injured cortex. Layer V is outlined. **b.** Quantification of percent area of IBA1 expression in the ipsilateral compared to contralateral cortex. **c.** Quantification of percent area of GFAP expression in the ipsilateral compared to contralateral cortex. N=6 per timepoint, 3-4 sections counted per animal. Error bars represent SEM. Scale bar, 200 µm.

### Supplemental Figure 7

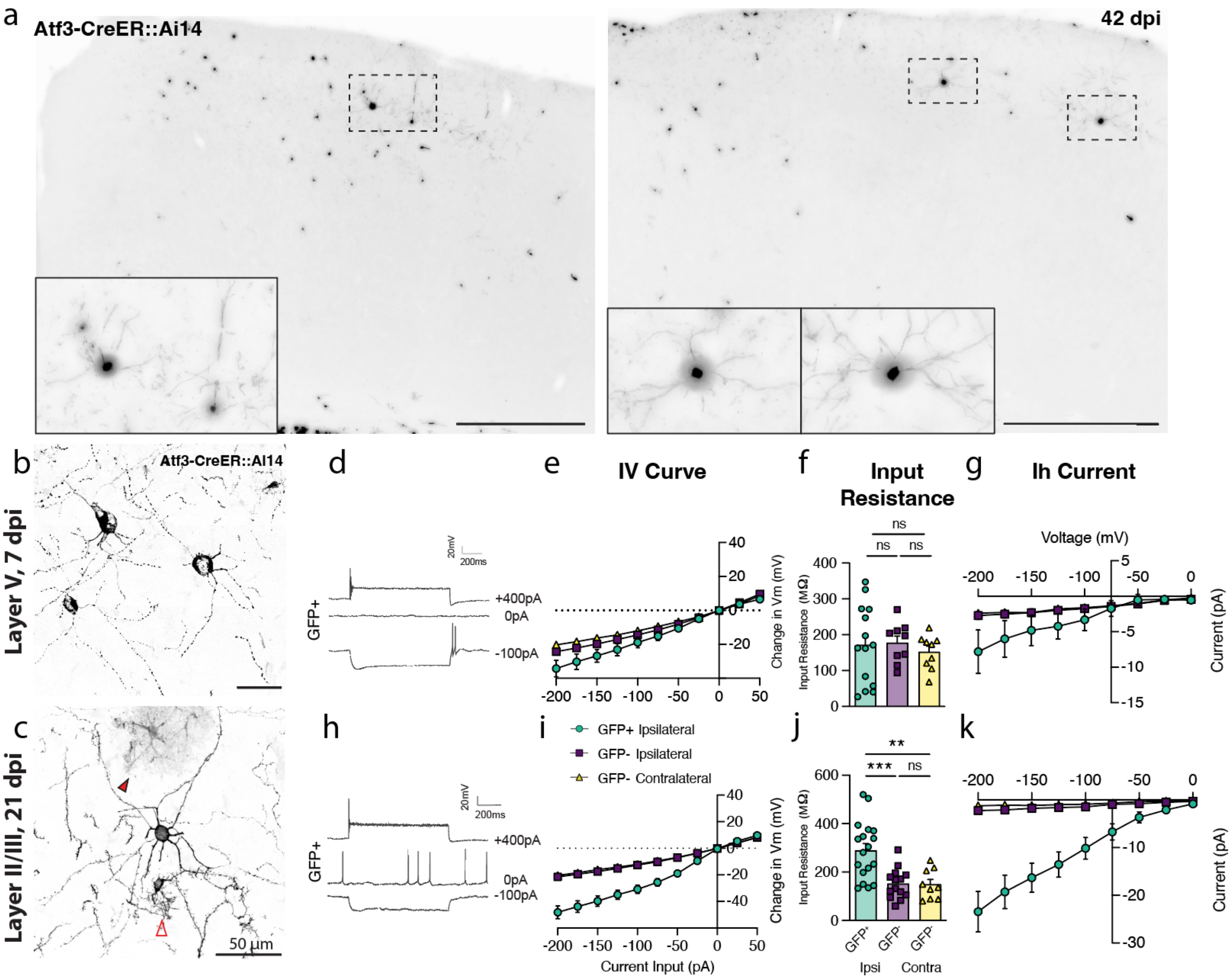

**Supplemental Figure 7. Layer II/III Atf3-expressing neuron survival and electrophysiological properties. a.** Low magnification images of the ipsilateral cortex of Atf3-CreER::Ai14 mTBI mice at 42 dpi showing surviving neurons and glia. Insets highlight examples of surviving layer II/III neurons. **b.** High magnification image highlighting degeneration of layer V Atf3-CreER::Ai14 neurons at 7 dpi. **c.** High magnification image highlighting an intact layer II/III Atf3-CreER::Ai14 neuron at 21 dpi. Image includes a Atf3-Ai14 microglion (bottom, open arrowhead) and astrocyte (top, closed arrowhead). **d.** Additional example traces of a layer V neuron at 7 dpi. Quantifications of **e.** IV curve, **f.** input resistance, **g.** Ih current in 7 dpi layer V neurons. **h.** Additional example traces of a layer II/III neuron at 21 dpi showing tonic firing. Quantifications of **i.** IV curve **j.** input resistance (** $p = 0.0018$, *** $p = 0.0005$ by Tukey's multiple comparisons test) **k.** Ih current in 21 dpi layer II/III neurons. For IV curve and Ih current graphs, points represent the average of all neurons per group, error bars represent SEM. For input resistance, each point represents one neuron recorded from N = 2-3 animals. Low magnification scale bars, 500 µm. High magnification scale bars, 50 µm.

### Supplemental Figure 8

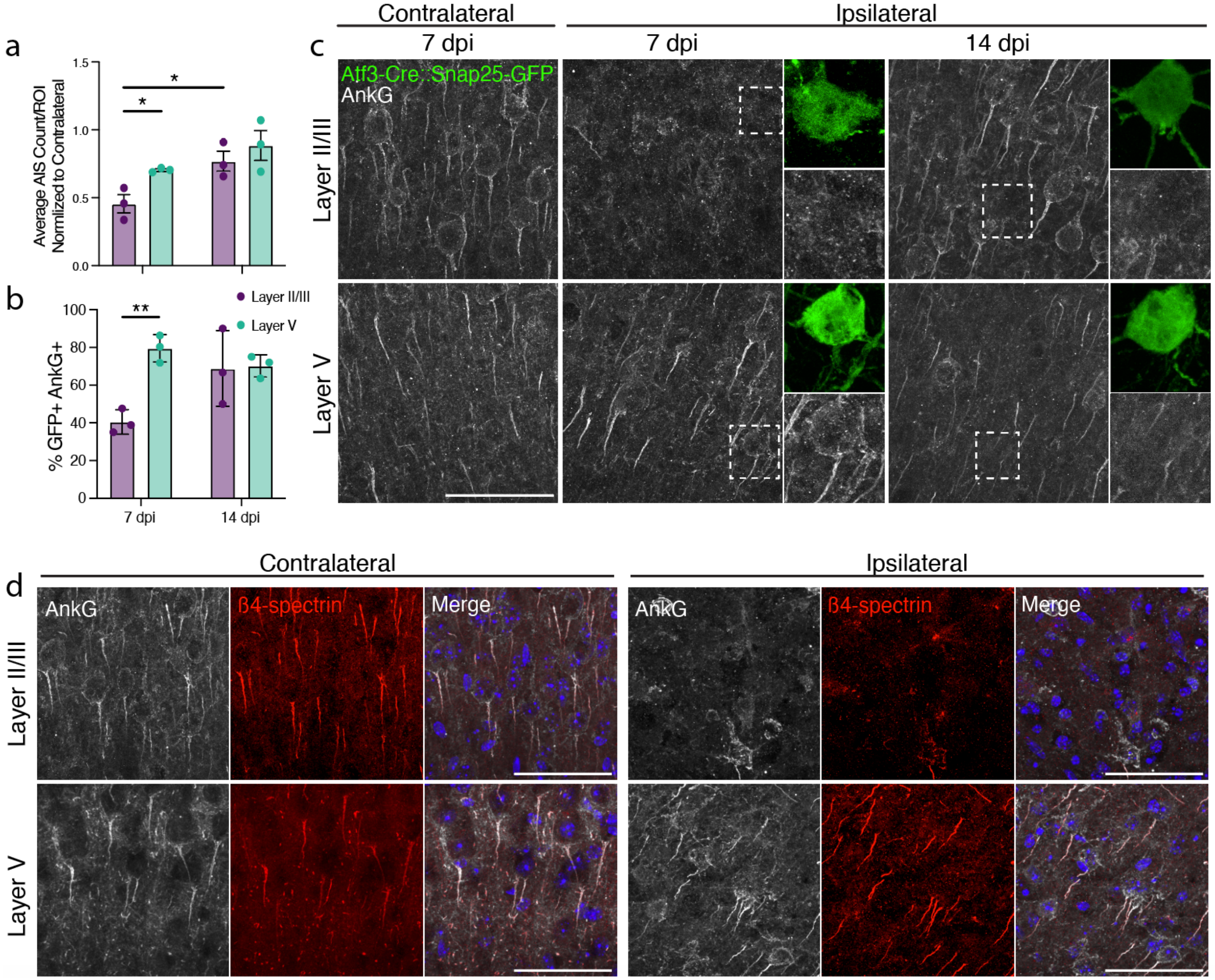

**Supplemental Figure 8. Axon initial segments are transiently lost in layer II/III neurons following mTBI.** **a.** Quantification of average number of AnkG+ axon initial segments in layer V and layer II/III at 7 dpi and 14 dpi showing a relative reduction in layer II/III at 7 dpi (* $p = 0.0224$ for layer V vs II/III, $p = 0.0353$ for 7 vs 14 dpi by unpaired t-test). **b.** Quantification of percent of GFP+ neurons in Atf3-GFP mice with an AnkG+ axon initial segment in layer V and layer II/III at 7dpi and 14 dpi showing a transient loss of axon initial segments in layer II/III neurons (** $p = 0.0023$ by unpaired t-test). **c.** Example 63X images of AnkG+ axon initial segments in the contralateral cortex at 7 dpi and in the ipsilateral cortex at 7 and 14 dpi in layer II/III and layer V. Insets highlight GFP+ neurons and their axon initial segments. **d.** Example high magnification images of AnkG and β4-spectrin in layer V and layer II/III in the contralateral and ipsilateral cortex at 7 dpi confirming loss of the axon initial segment and not AnkG immunoreactivity. Merge includes DAPI expression. For **a** and **b**, each point represents the average of 3-4 ROIs per 3-4 sections per animal. Scale bars, 50 µm.

### Supplemental Figure 9

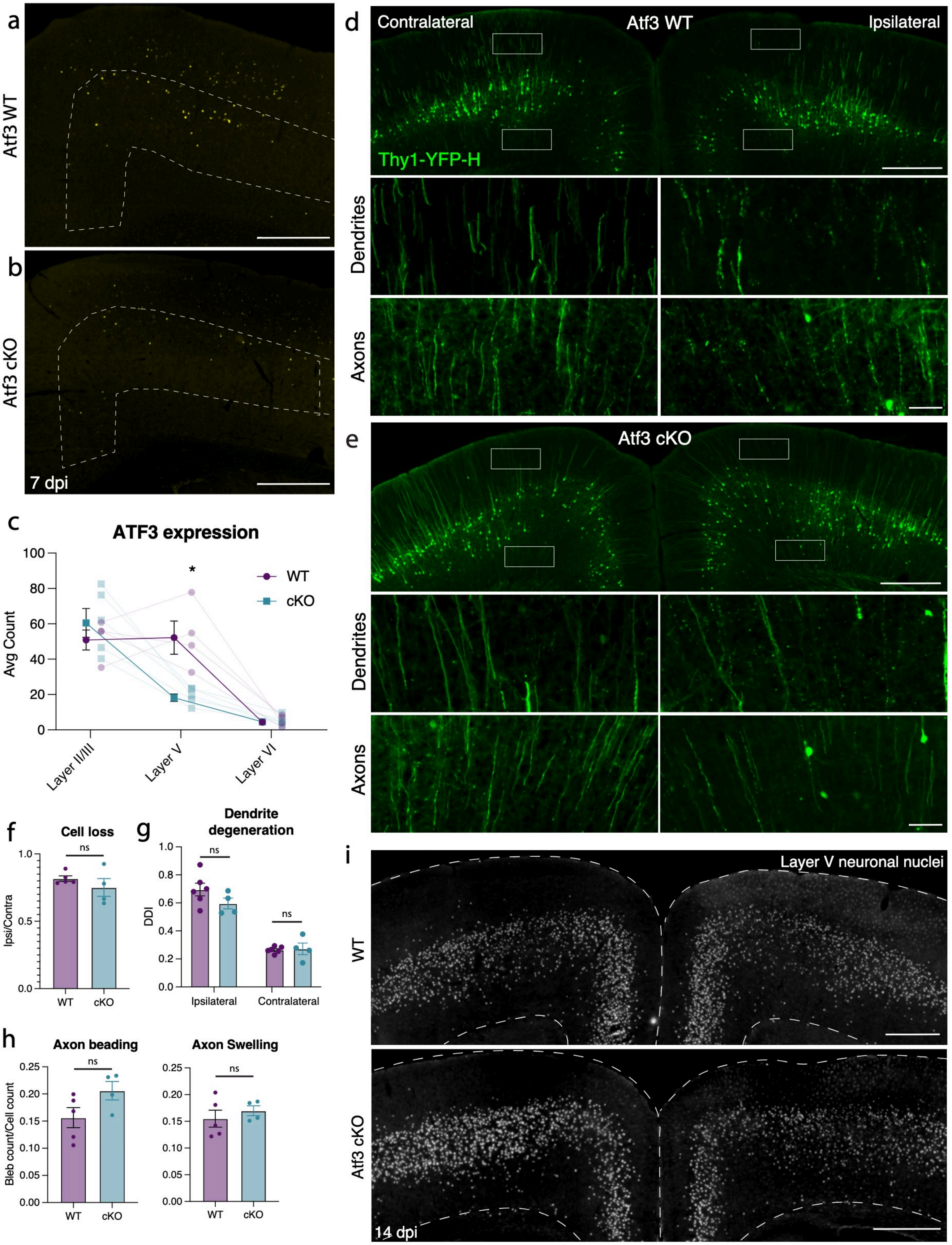

**Supplemental Figure 9. Deletion of Atf3 in layer V neurons is not sufficient to prevent neuron degeneration. a.** ATF3 expression in ipsilateral cortex at 7 dpi in a wildtype (WT) mouse. Layer V is outlined. **b.** ATF3 expression in the ipsilateral cortex of an Atf3 cKO mouse highlighting a reduction of Atf3 in layer V (outlined). Remaining cells are likely neurons outside of the Rbp4-Cre lineage or ATF3-expressing glia. **c.** Quantification of ATF3 protein expression by cortical layer validating layer V-specific knockout in Atf3 cKO animals. Remaining expression is likely non-neuronal. Average and SEM are shown for each cortical layer with more transparent points representing individual animals and the average of 3-4 sections (* $p = 0.033$ by t-test between genotypes in layer V). **d.** Thy1-YFP expression in a WT animal with insets highlighting dendrites and axons in the ipsilateral and contralateral cortices. **e.** Thy1-YFP expression in an Atf3 cKO animal with insets highlighting dendrites and axons on the ipsilateral and contralateral cortices. **f.** Quantification of nuclei in the ipsilateral cortex normalized to contralateral cortex at 14 dpi based on such images as in i showing no change in Atf3 cKO. **g.** Dendrite degeneration quantifications from Thy1-YFP animals (examples shown in d,e) revealing no change in Atf3 cKO. **h.** Axon beading and swelling quantifications from Thy1-YFP animals (examples shown in **d,e**) revealing no change in Atf3 cKO. **i.** Cre-dependent labeling of nuclei in WT and Atf3 cKO animals at 14 dpi. ns: not significant. Scale bars, 500 µm.

### Supplemental Figure 10

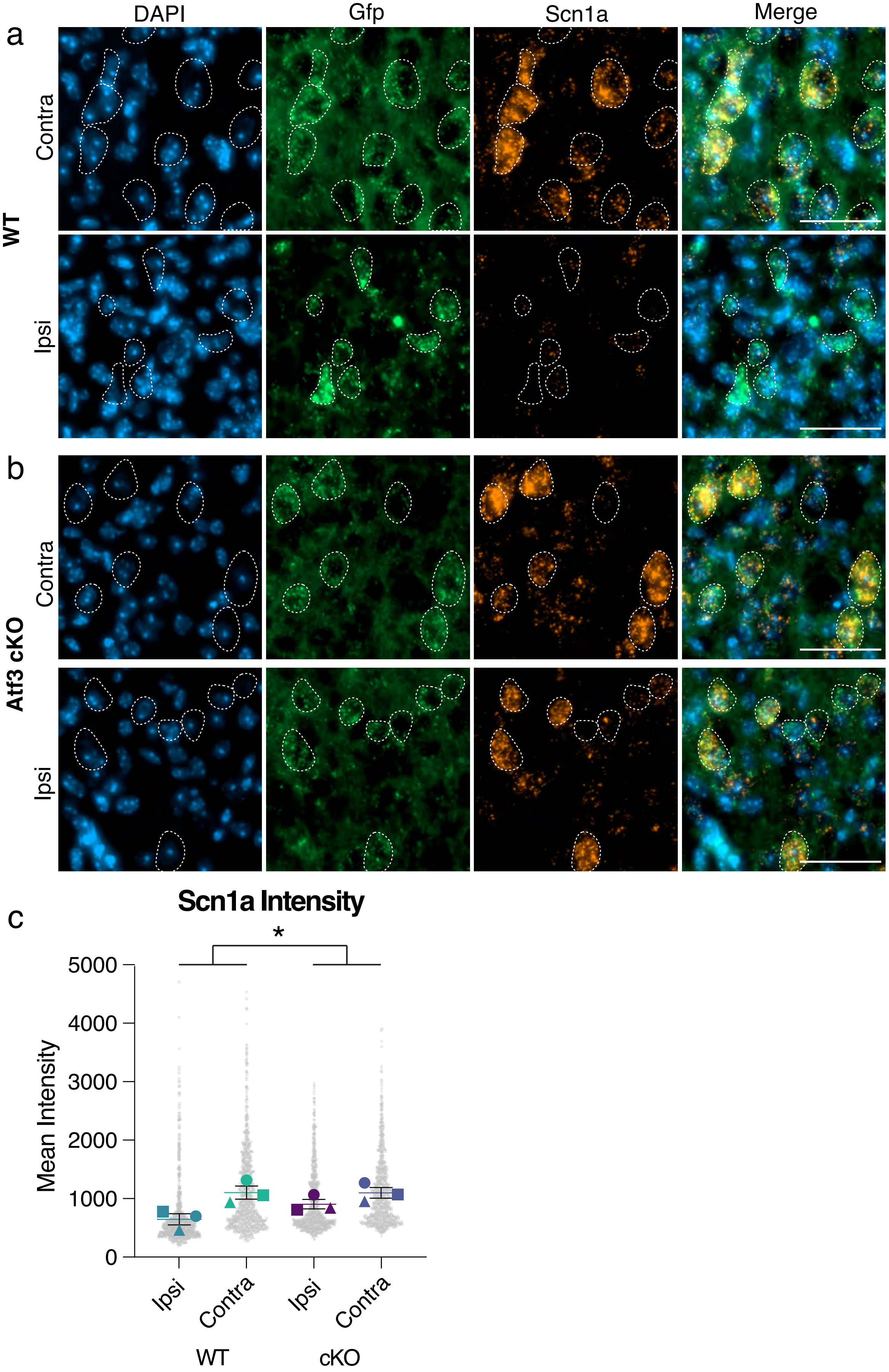

**Supplemental Figure 10. mTBI-induced *Scn1a* downregulation is *Atf3*-dependent. a.** Multiplexed *in situ* hybridization showing expression of DAPI, *Gfp*, and *Scn1a* in layer V of a WT mouse (Gfp induced by Rbp4-Cre::Sun1-GFP reporter for specific measurement in layer V neurons) revealing downregulation of *Scn1a* in *Gfp*+ neurons (outlined) at 7 dpi. **b.** Multiplexed *in situ* hybridization showing expression of DAPI, *Gfp*, and *Scn1a* in layer V of an Atf3 cKO mouse (Gfp induced by Rbp4-Cre::Sun1-GFP reporter for specific measurement in layer V neurons) revealing prevention of *Scn1a* downregulation in *Gfp*+ neurons (outlined) at 7 dpi. **c.** Quantification of mean intensity of *Scn1a* in layer V neurons at 7 dpi showing prevention of *Scn1a* downregulation in Atf3 cKO neurons (interaction of ipsi vs contra in WT vs cKO * $p = 0.01$ by linear mixed effect modeling). Average per animal and value per cell are displayed. Each shape represents one animal. Scale bars, 50 µm.

### Supplemental Figure 11

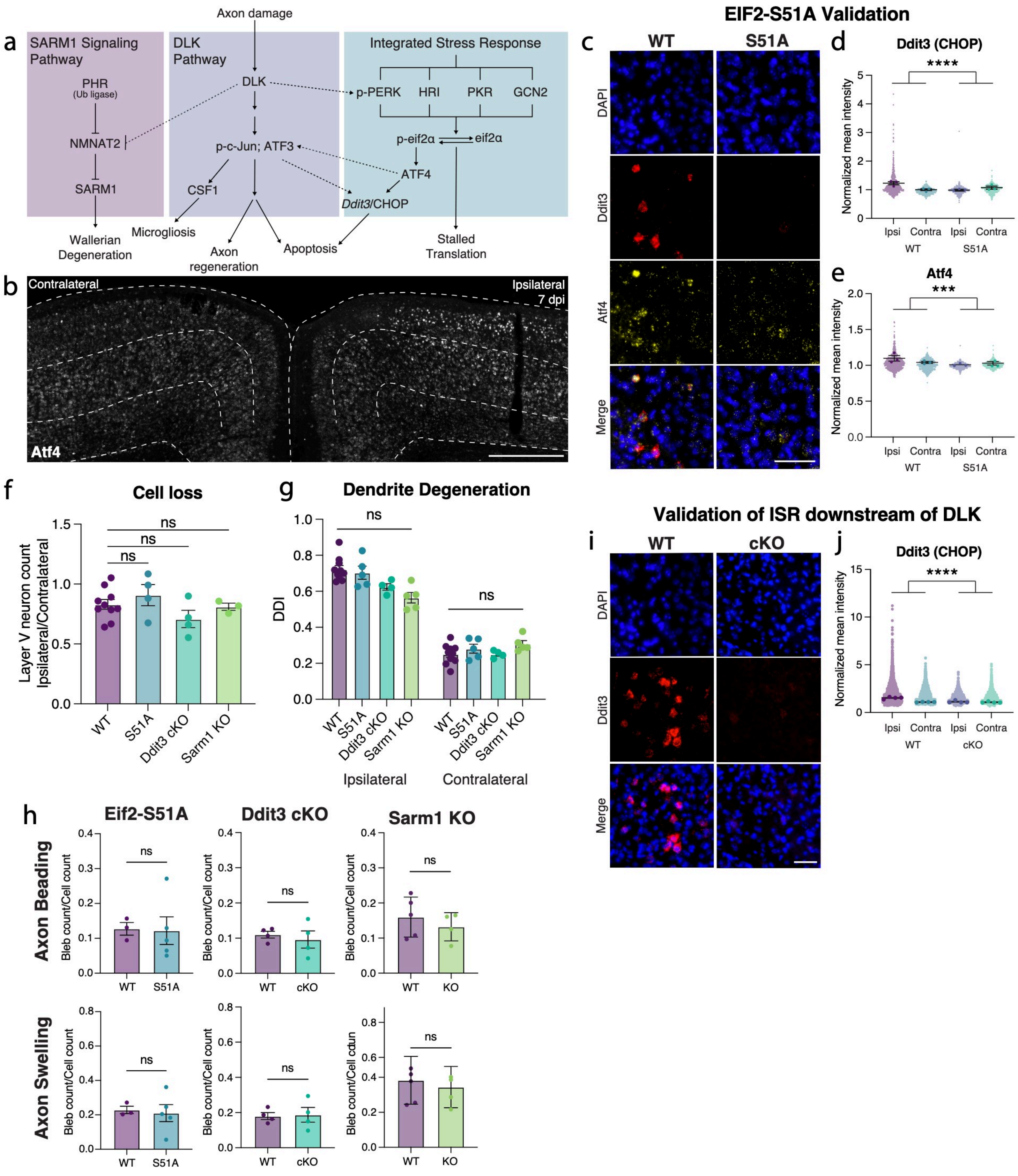

**Supplemental Figure 11. Stress response pathways downstream of DLK are not sufficient to induce layer V neuron death. a.** A diagram summarizing the interactions between the DLK, SARM1, and integrated stress response pathways and their major effectors. **b.** *In situ* hybridization of Atf4, the major effector of the integrated stress response, in the WT mTBI cortex at 7 dpi showing increased mRNA expression in ipsilateral layer V and layer II/III. Layer V is outlined. **c.** Validation of ISR knockdown in the EIF2-S51A mouse, showing in situ hybridization of Atf4 and Ddit3 in ipsilateral layer V of WT and S51A mice. **d.** Quantification of Ddit3 (protein name CHOP) *in situ* hybridization in ipsilateral and contralateral layer V of WT and S51A animals (interaction of ipsi vs contra in WT vs S51A **** $p = 1.54e-09$ by linear mixed effect modeling). **e.** Quantification of Atf4 in situ hybridization in ipsilateral and contralateral layer V of WT and S51A animals (interaction of ipsi vs contra in WT vs S51A *** $p = 0.0026$ by linear mixed effect modeling). **f.** Quantification of cell loss in layer V of all knockout lines at 14 dpi measured as the ratio of ipsilateral to contralateral cell count, showing that cell loss is insignificantly improved by each manipulation. For all cKO mice, Cre-dependent Sun1-GFP was counted. For KO animals, Thy1-YFP cell bodies were counted. N= 5 WT Thy1-YFP and N= 5 WT cKO were pooled in the WT condition. **g.** Quantification of dendrite degeneration in all knockout lines at 7 dpi showing insignificant changes compared to wildtype. Shapes represent littermates from each strain. **h.** Quantification of axon pathology in the ipsilateral cortex of all knockout lines, measured as axon beading (fragments with area < 10 µm2) or axon swellings (fragments with area > 10 µm2), showing insignificant changes compared to wildtype. All knockouts are compared to littermate controls. **i.** Validation that the ISR is downstream of Dlk showing in situ hybridization of Atf4 and Ddit3 in ipsilateral layer V of Dlk WT and Dlk cKO mice. **j.** Quantification of Ddit3 (protein name CHOP) in situ hybridization in ipsilateral and contralateral layer V of Dlk WT and Dlk cKO animals (ipsi vs contra in WT vs cKO **** $p = 5.47e-09$ by linear mixed effect modeling). **k.** Quantification of Atf4 in situ hybridization in ipsilateral and contralateral layer V of Dlk WT and Dlk cKO animals. For **d, e, j, k**, average per animal and value per cell are displayed. For **f-h**, each point represents the average of 3-4 sections per animal. Low magnification scale bars, 500 µm. High magnification scale bar, 50 µm.

### Supplemental Figure 12

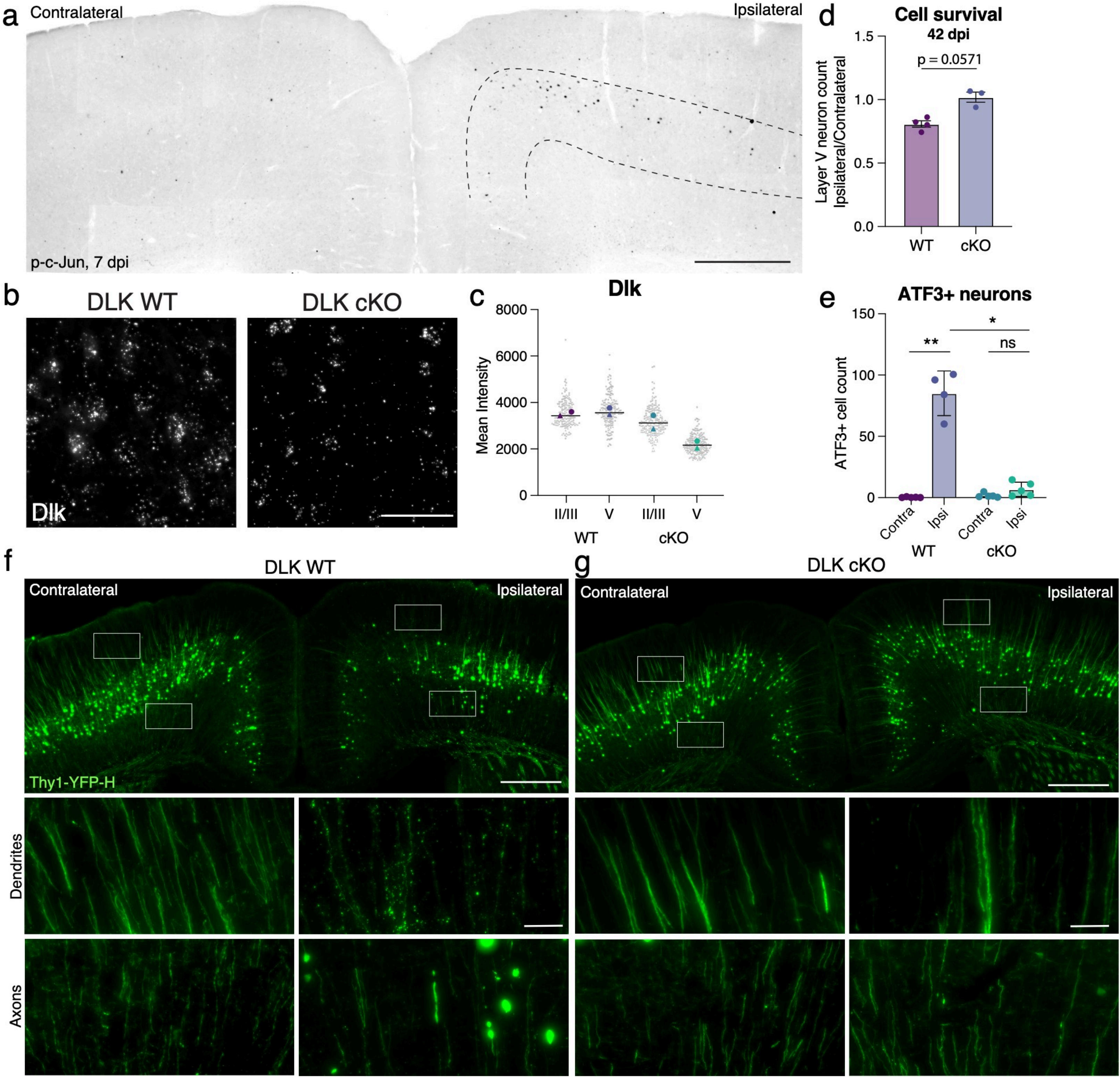

**Supplemental Figure 12. Dlk cKO leads to a reduction in layer V Dlk mRNA and is neuroprotective. a.** Low magnification image of p-c-Jun in the ipsilateral and contralateral cortices of a WT mouse at 7 dpi showing higher expression in ipsilateral layer V (outlined) and no expression in the contralateral cortex. **b.** High magnification images of ISH for full-length Dlk mRNA in layer V in WT and cKO animals showing a reduction in Dlk cKO. The full length probe still detects mRNA sequence in the cKO animals, with lower intensity signal reflecting cKO. **c.** Validation of DLK cKO by in situ hybridization. Quantification of mean intensity of Dlk mRNA in layer II/III and layer V of WT and cKO animals showing a layer V-specific reduction in cKO animals. Average per animal and value per cell are displayed. Each shape represents one animal. Quantifications from N=2 animals and 2 sections per animal. **d**. The rescue of layer V neuron loss was maintained in Dlk cKO neurons at 42 dpi ($p = 0.0571$ by t-test). **e.** Quantification of the number of ATF3+ neurons in layer V in WT and Dlk cKO animals confirming that ATF3 activation is Dlk-dependent. This was measured in Dlk cKO animals crossed to the Sun1-GFP reporter such that only neurons in which Dlk was deleted were counted (* $p = 0.0138$, ** $p = 0.0097$ by Brown-Forsythe and Welch test). **f.** Thy1-YFP expression in a WT animal with insets highlighting dendrites and axons on the ipsilateral and contralateral cortices. **g.** Thy1-YFP expression in a Dlk cKO animal with insets highlighting dendrites and axons on the ipsilateral and contralateral cortices. Low magnification scale bar, 500 µm. High magnification scale bar, 50 µm.

### Supplemental Figure 13

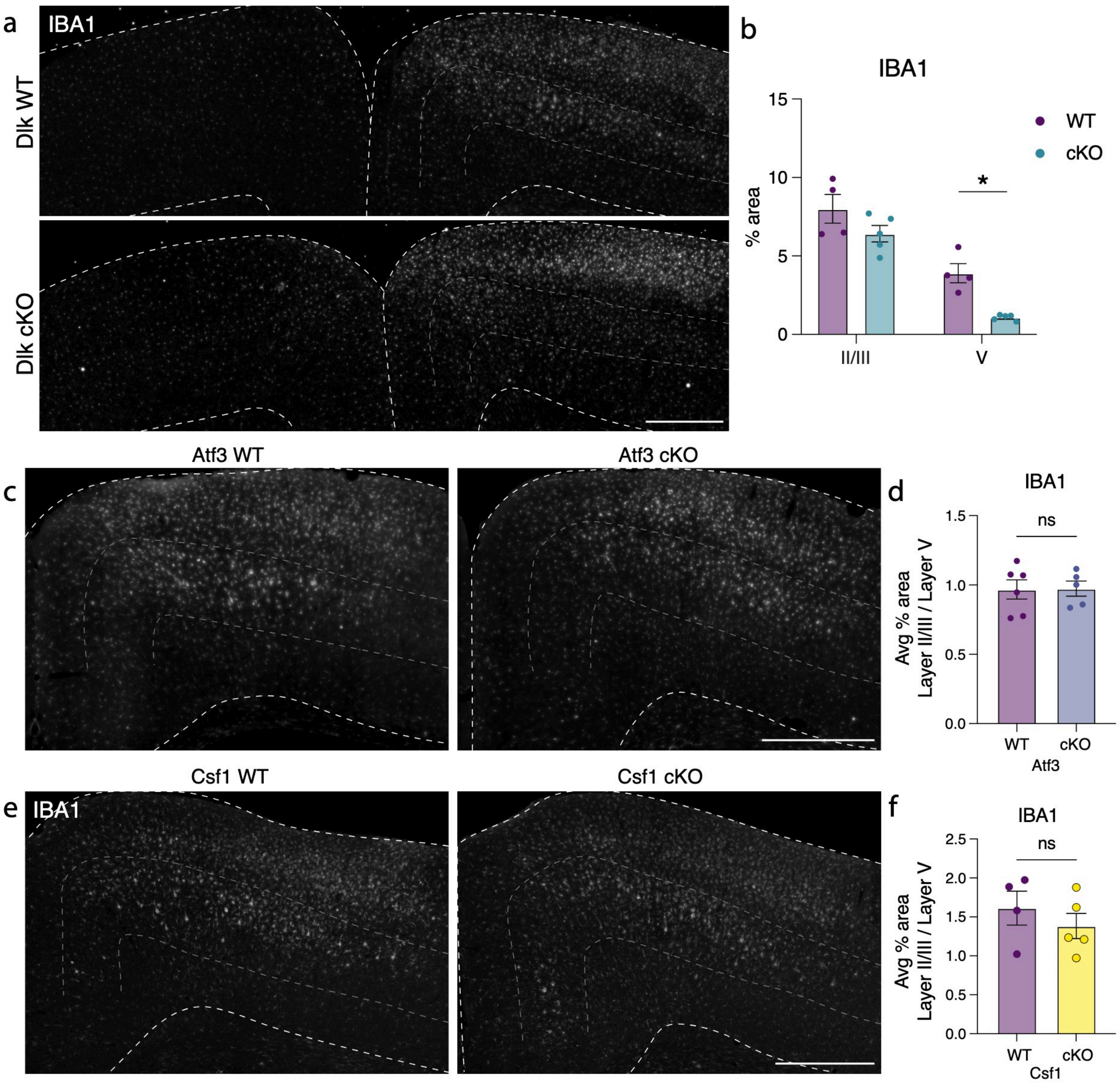

**Supplemental Figure 13. DLK deletion leads to a decrease in cortical microgliosis in a Csf1- and Atf3-independent manner. a.** IBA1 immunostaining in a DLK WT and Dlk cKO cortex showing that microgliosis is reduced in layer V, where DLK is deleted. Layer V is outlined. **b.** Quantification of percent area of IBA1 immunostaining in layer II/III compared to layer V in WT and cKO animals (* p = 0.0190 by unpaired t-test). **c.** Immunostaining of IBA1 in the ipsilateral cortex of Atf3 WT and cKO animals at 7 dpi showing no effect on microgliosis. Laye r V is outlined. **d.** Quantification of IBA1 percent area in Atf3 WT and cKO cortices, shown as layer II/III percent area over layer V percent area. **e.** Immunostaining of IBA1 in the ipsilateral cortex of Csf1 WT and cKO animals at 7 dpi showing that cortical microgliosis after mTBI is not Csf1-dependent. Layer V is outlined. **f.** Quantification of IBA1 percent area in Csf1 WT and cKO cortices, shown as layer II/III percent area over layer V percent area. ns: not significant. Scale bars, 500 µm.
